# Genetic diversity in two *Plasmodium vivax* protein ligands for reticulocyte invasion

**DOI:** 10.1101/328757

**Authors:** Camille Roesch, Jean Popovici, Sophalai Bin, Vorleak Run, Saorin Kim, Stéphanie Ramboarina, Emma Rakotomalala, Rado Lalaina Rakotoarison, Tsikiniaina Rasoloharimanana, Zo Andriamanantena, Anuj Kumar, Micheline Guillotte-Blisnick, Christèle Huon, David Serre, Chetan E. Chitnis, Inès Vigan-Womas, Didier Menard

**Affiliations:** Malaria Molecular Epidemiology Unit, Institut Pasteur in Cambodia, Phnom Penh, Cambodia; Immunology of Infectious Diseases Unit, Institut Pasteur de Madagascar, Antananarivo, Madagascar; International Centre for Genetic Engineering and Biotechnology (ICGEB), New Delhi, India; Malaria Parasite Biology and Vaccines Unit, Institut Pasteur, Paris, France; Institute for Genome Sciences, University of Maryland, Baltimore, Maryland, USA

## Abstract

The interaction between *Plasmodium vivax* Duffy binding protein (PvDBP) and Duffy antigen receptor for chemokines (DARC) has been described as critical for the invasion of human reticulocytes, although increasing reports of *P. vivax* infections in Duffy-negative individuals questions its unique role. To investigate the genetic diversity of the two main protein ligands for reticulocyte invasion, PvDBP and *P. vivax* Erythrocyte Binding Protein (PvEBP), we analyzed 458 isolates collected in Cambodia and Madagascar. First, we observed a high proportion of isolates with multiple copies PvEBP from Madagascar (56%) where Duffy negative and positive individuals coexist compared to Cambodia (19%) where Duffy-negative population is virtually absent. Whether the gene amplification observed is responsible for alternate invasion pathways remains to be tested. Second, we found that the *PvEBP* gene was less diverse than *PvDBP* gene (12 *vs.* 33 alleles) but provided evidence for an excess of nonsynonymous mutations with the complete absence of synonymous mutations. This finding reveals that PvEBP is under strong diversifying selection, and confirms the importance of this protein ligand in the invasion process of the human reticulocytes and as a target of acquired immunity. These observations highlight how genomic changes in parasite ligands improve the fitness of *P. vivax* isolates in the face of immune pressure and receptor polymorphisms.

## Author summary

Until recently, *P. vivax* was thought to infect only Duffy positive individuals, due to its dependence on binding the Duffy blood group antigen as a receptor for reticulocyte invasion and to be absent from parts of Africa where the Duffy-negative phenotype is highly frequent. However, a number of recent studies from across sub-Saharan Africa have reported *P. vivax* infections in Duffy-negative individuals. Invasion into Duffy-positive reticulocytes is mediated by the *P. vivax* Duffy binding protein (PvDBP). The mechanism for invasion into Duffy-negative reticulocytes is not known. A homologue of PvDBP, namely, *P. vivax* erythrocyte binding protein (PvEBP), has been recently identified but its role in Duffy independent invasion is not clearly defined. Here, we provide unique insights into the roles of these two key ligands by studying the genetic diversity of *P. vivax* isolates collected from Cambodia, where all individuals are Duffy positive, and Madagascar where both Duffy-positive and Duffy-negative individuals coexists. Our data suggest that PvEBP may play an important functional role in invasion into Duffy-negative reticulocytes. PvEBP appears to be a target of naturally acquired antibody responses following natural exposure to *P. vivax* infection and such as a consequence an important vaccine candidate, together with PvDBP.

## Introduction

*Plasmodium vivax* is a predominant cause of malaria outside Africa, which causes significant morbidity (estimate of 8.5 million cases in 2016) and places an enormous economic burden on many resource poor countries [1]. Until recently, vivax malaria was considered a benign infection compared to *P. falciparum*, although clinical episodes and regular recurrent infections cause significant morbidity as well as impact on the economy in endemic areas [2]. Moreover, *P. vivax* infections can sometimes lead to severe life-threatening pathologies [3].

Previous data from malaria therapy, which was used extensively for over two decades (1920-1940) for treatment of neurosyphilis (e.g. paralysis of the insane), as well as experimental infections of volunteers have demonstrated that individuals of African origin were naturally resistant to *P. vivax* infection [4–6]. Thereafter, following identification of the Duffy blood group [7], it was shown that the Duffy blood group antigen (Fy^a^ or Fy^b^) was not expressed on red blood cells (RBCs) of individuals of African origin (Duffy-negative) [8, 9]. Seminal works with controlled experimental infections of volunteers through sporozoite challenge and *in vitro* invasion studies using *P. knowlesi* as a model subsequently established the paradigm that the Duffy antigen is required for reticulocyte invasion by *P. vivax* [9–11]. Consequently, vivax malaria was long thought to be absent from parts of Africa where the Duffy-negative phenotype is highly frequent [12].

Host cell invasion by *Plasmodium* merozoites is a complex, multi-step process that involves multiple interactions between erythrocyte receptors and ligands on merozoites. Unlike *P. falciparum* merozoites, which can use several erythrocyte receptors for invasion, it was thought, until recently, that invasion of human reticulocytes by *P. vivax* is completely dependent on the interaction between the *P. vivax* Duffy Binding Protein (PvDBP) and the erythrocyte Duffy antigen receptor for chemokines (DARC) [13, 14]. However, over the last decade, a growing body of studies have reported PCR-positive vivax malaria cases in Duffy-negative individuals around the world, specifically across Africa (review in [12]) and South America [15, 16]. These frequent observations raise the emerging issue of *P. vivax* infection in Duffy-negative populations, and the possibility of alternative invasion mechanism(s). At present, we do not know whether these clinical reports are common and were previously undetected or if they are due to the emergence and spread of specific *P. vivax* strains that use alternative Duffy-independent pathways to invade Duffy-negative reticulocytes. Whatever the reason for the increasing frequency of observations of *P. vivax* in Duffy-negative populations, it is a cause for concern and demands attention. Recent whole genome sequencing studies of monkey-adapted *P. vivax* strains and field isolates (all isolates were collected from Duffy-positive patients) have revealed that the *PvDBP* gene was duplicated in multiple *P. vivax* isolates, particularly at high prevalence in Madagascar, a setting where Duffy-positive and Duffy-negative individuals coexist [17]. Initially, this epidemiological pattern suggested that the duplication of this gene was likely associated with the capability of the parasite to overcome the barrier of Duffy negativity. Since then, this hypothesis has been challenged by Hostetler *et al.* [18] who found that *PvDBP* gene duplications were widespread even in malaria endemic areas in Southeast Asia where Duffy-negativity is not present. Another recent study [19] reported evidence of *PvDBP* gene amplification (3 and 8 copies) in two Duffy-negative Ethiopian isolates. In addition, sequence data generated from a *P. vivax* field isolate (C127 isolate from Cambodia), which used reconstruction of long reads without relying on the reference genome (e.g. the monkey-adapted Salvador I strain), identified 792 predicted genes [20]. Among them, two contigs contained predicted protein coding genes similar to known *Plasmodium* red blood cell invasion proteins. One of these genes harbored all the hallmarks of a *Plasmodium* erythrocyte binding protein, including conserved Duffy-binding like and C-terminal cysteine-rich domains. Further analysis showed that this gene, which is present in most of studied *P. vivax* genomes, clustered separately from all known *Plasmodium erythrocyte-binding protein* genes [20]. Further functional investigations demonstrated that the recombinant PvEBP (derived from the C127 *PvEBP* allele) bound preferentially to immature (CD71^high^) and Duffy-positive reticulocytes [21]. A minimal binding was observed with Duffy-negative reticulocytes, and no binding was observed with mature red blood cells or normocytes. PvDBP and PvEBP were clearly shown to be antigenically distinct. These findings were slightly modulated in another study that reported that the region II of PvEBP expressed in COS-7 cells bound both Duffy-positive and Duffy-negative erythrocytes although at low frequency suggesting that PvEBP may be a ligand for invasion of Duffy-negative reticulocytes [19].

To gain insight into the natural genetic diversity and polymorphisms in the two main protein ligands for reticulocyte invasion (*PvDBP* and *PvEBP*), we collected and analyzed *P. vivax* isolates from two distinct settings: in Cambodia, where the vast majority of individuals are Duffy-positive[2], and Madagascar where an admixture of Duffy-positive and Duffy-negative people coexists [22].

## Results

### Duffy genotyping

Duffy genotyping data were available for 174 samples and all samples were genotyped as Duffy-positive (T-33C substitution: Cambodia, N=119, 100% T/T and Madagascar, N=55, 44% T/T and 56%T/C) (Table 1).

**Table 1.** Number of samples from Cambodia and Madagascar collected before (D0) and after treatment in case of recurrence (Dx) with available data for *PvDBP* and *PvEBP* (SNP and CNV) and Duffy genotyping.

| No. of samples with available data for | Day of collection | Country | Year |  |  |  |  |  |  |  |  | Total |
| --- | --- | --- | --- | --- | --- | --- | --- | --- | --- | --- | --- | --- |
|  |  |  | 2003 | 2004 | 2011 | 2012 | 2013 | 2014 | 2015 | 2016 | 2017 |  |
| SNPs<br>*PvDBP* | D0 | Cambodia |  |  |  |  | 80 | 18 | 46 | 9 |  | 153 |
|  | Dx |  |  |  |  |  |  |  |  |  |  | 0 |
|  | D0 | Madagascar |  |  |  |  |  |  | 25 | 57 | 10 | 92 |
| SNPs<br>*PvEBP* | D0 | Cambodia |  |  |  |  | 87 | 17 | 36 | 10 |  | 150 |
|  | Dx |  |  |  |  |  |  |  |  |  |  | 0 |
|  | D0 | Madagascar |  |  |  |  |  |  | 27 | 39 | 3 | 69 |
| CNVs<br>*PvDBP* | D0 | Cambodia | 2 | 46 | 51 | 48 | 73 | 94 | 41 | 17 | 20 | 392 |
|  | Dx |  |  |  |  |  |  | 14 |  |  |  | 14 |
|  | D0 | Madagascar |  |  |  |  |  |  | 25 | 25 | 16 | 66 |
| CNVs<br>*PvEBP* | D0 | Cambodia |  |  |  |  | 59 | 42 | 47 | 16 | 20 | 184 |
|  | Dx |  |  |  |  |  |  | 5 |  |  |  | 5 |
|  | D0 | Madagascar |  |  |  |  |  |  | 23 | 27 | 16 | 66 |
| Duffy genotyping | D0 | Cambodia |  |  |  |  | 13 | 17 | 53 | 16 | 20 | 119 |
|  | Dx | Madagascar |  |  |  |  |  |  | 27 | 24 | 4 | 55 |
| **Total number** | **D0** | **Cambodia** | **2** | **46** | **51** | **48** | **92** | **106** | **54** | **18** | **20** | **437** |
|  | **Dx** |  |  |  |  |  |  | **19** |  |  |  | **19** |
|  | **D0** | **Madagascar** |  |  |  |  |  |  | **44** | **67** | **18** | **129** |

### *PvDBP* gene copy number variation (CNV)

*PvDBP* CNV was assessed in 458 *P. vivax* isolates collected in Cambodia (N=392) and Madagascar (N=66) (Table 1). *PvDBP* CNV ranged from 1 to 6 copies. No significant differences were observed between *P. vivax* isolates from both countries in the median gene copy number (1.31 and 1.30 for Madagascar and Cambodia, respectively, P=0.21, Mann Whitney test) or in the proportion of isolates carrying multiple copies *PvDBP* (45.5% for Madagascar and 37.5% for Cambodia, P=0.22, Fisher’s exact test). These data confirm that *PvDBP* amplification is a common event across isolates from Cambodia and Madagascar (Fig 1).

**Fig 1.**
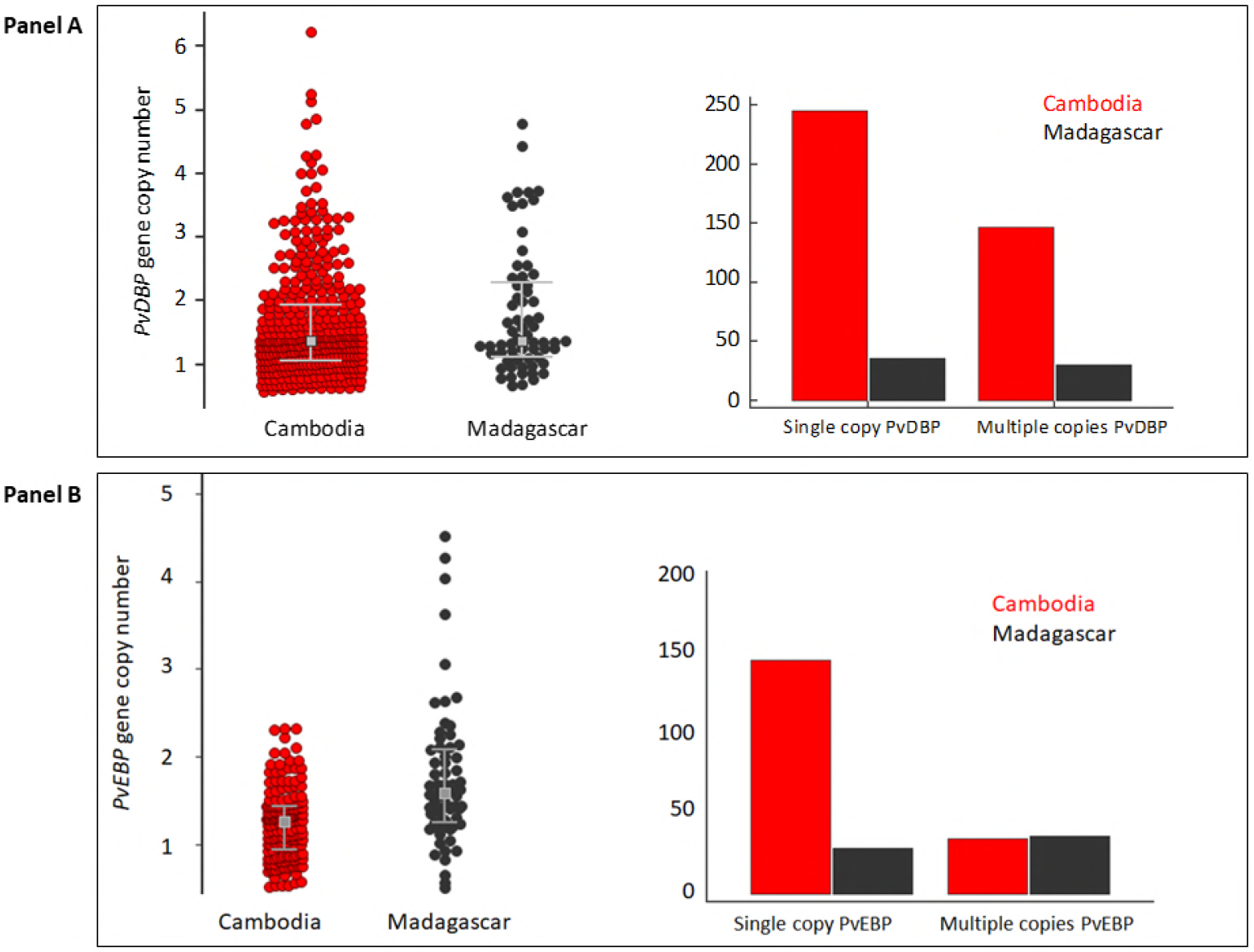
Distribution of *PvDBP* and *PvEBP* gene copy number in isolates from Cambodia and Madagascar (Panel A and B, left side). Number of isolates from Cambodia and Madagascar with single or multiple copies *PvDBP* and *PvEBP* genes (Panel A and B, right side). The grey squares represent the medians and whiskers the IQRs.

Further analysis showed that the proportion of isolates with multiple copies of *PvDBP* increased over time in Cambodia: from 16.7% (8/48) in 2003-2004 to 39.8% (142/357) in 2011-2017 (P=0.001, Fisher’s exact test). This trend was significant for samples collected from western provinces (Battambang, Pursat, Pailin, Kampot) (15.2% *vs.* 36.6%, P=0.02, Fisher’s exact test). The proportion of isolates with multiple copies of *PvDBP* and the median gene copy numbers for *PvDBP* were similar between paired isolates collected before a standard 3-day course of chloroquine (30 mg/kg) (D0) and at the day of recurrence (Dx) occurring during the 2-months follow up in patients relocated in a non-transmission area (D0: 38.5%, 5/13, median *PvDBP* copy number= 1.4, IQR: 1.1-2.4 vs. Dx: 23.1%, 3/13, median *PvDBP* copy number=1.0, IQR: 1.0-1.5, P=0.67, Fisher’s exact test and P=0.40, Mann Whitney test, respectively).

More surprisingly, we observed in Malagasy samples that the proportion of isolates with *PvDBP* amplification or the median copy of *PvDBP* gene was significantly higher in homozygous Duffy-positive individuals (T/T) compared to heterozygous Duffy-positive individuals (T/C) who are supposed to express less DARC antigen on the surface of their reticulocytes: T/T: 59.1%, 13/22, median *PvDBP* copy number= 2.2, IQR: 1.0-3.0 *vs.* T/C: 27.6%, 8/29, median *PvDBP* copy number=2.0, IQR: 0.9-1.5, P=0.04 (Fisher’s exact test and P=0.01, Mann Whitney test, respectively).

### *PvDBP* gene amplification occurs across multiple alleles

To determine whether *PvDBP* gene amplification was restricted to specific alleles, *PvDBPII* sequences (from codon 184 to codon 468) were determined among 153 Cambodian and 92 Malagasy *P. vivax* isolates. As already described, *PvDBP* sequences were found to be highly polymorphic and multiple alleles were observed both in Cambodia (21 alleles) and Madagascar (15 alleles) (Fig 2 and S1 Table).

**Fig 2.**
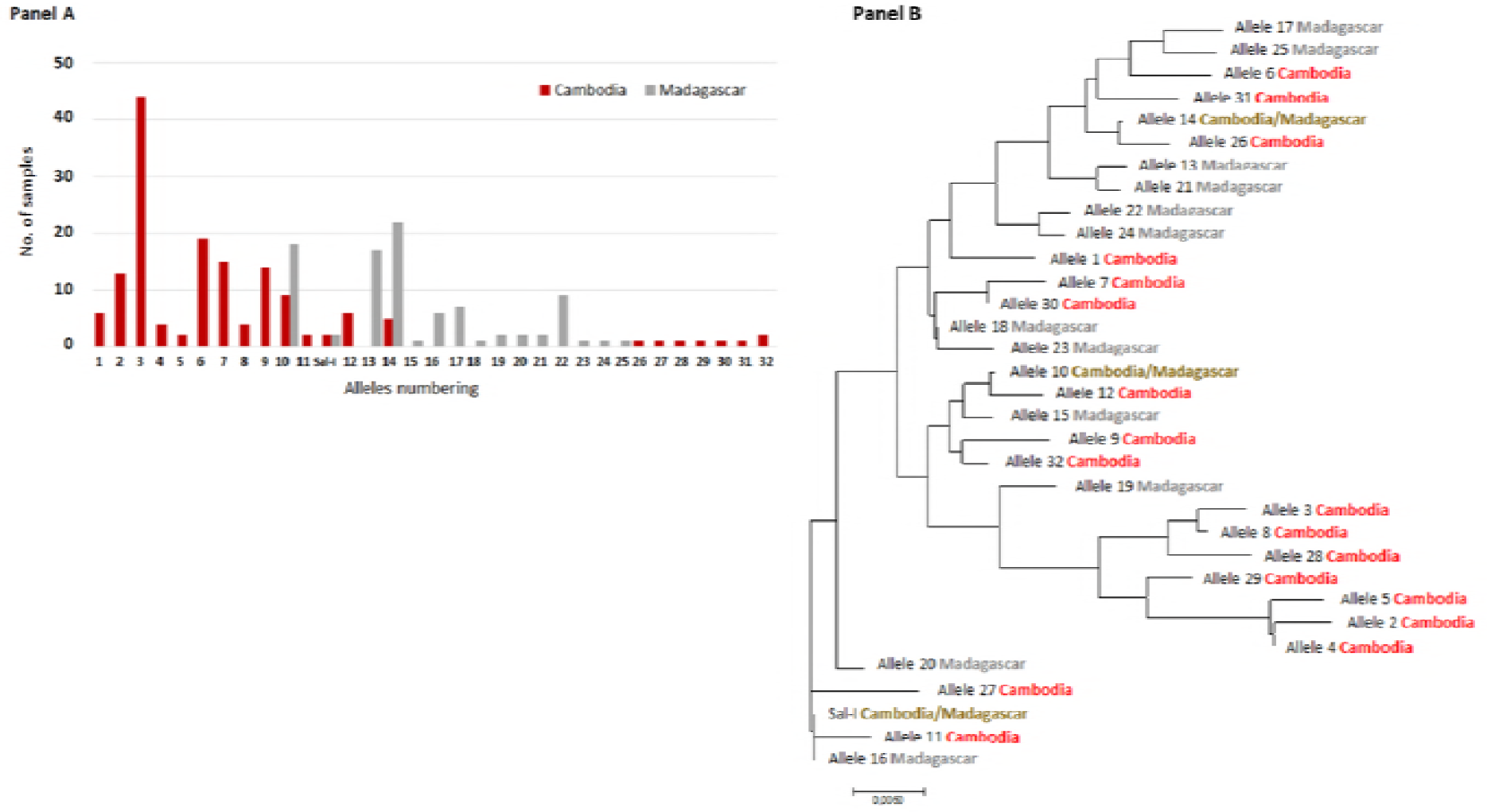
Distribution of *PvDBP* alleles observed in *P. vivax* isolates collected in Cambodia and Madagascar (Panel A). Phylogenetic tree inferred using the Neighbor-Joining method (Panel B). Each allele was numbered 1-32 (see S1 Table). The evolutionary distances were computed using the Poisson correction method [23] and are in the units of the number of amino acid substitutions per site. The analysis involved 33 amino acid sequences. Evolutionary analyses were conducted in MEGA7 [24].

Numerous SNPs already described were observed [25–37]. Four new SNPs (one silent and three non-synonymous) were detected (K260E, P450R, V453V and P475R). Among the 23 SNPs, some were solely observed in Madagascar (I419M, V453V and I464I) and in Cambodia (K260E, F306L, I367T, N375D, R378R, S398T, T404R, P450R, P475A/R and Q486E).

SNPs along with the location of binding residues for DARC, identified by previous studies [38–40], were mapped on to the 3D structure of PvDBPII. Interestingly, majority of the SNPs were found to be distal from the DARC binding residues, and located on the opposite surface compared to the binding residues for DARC (Fig 3, Panel A).

**Fig 3.**
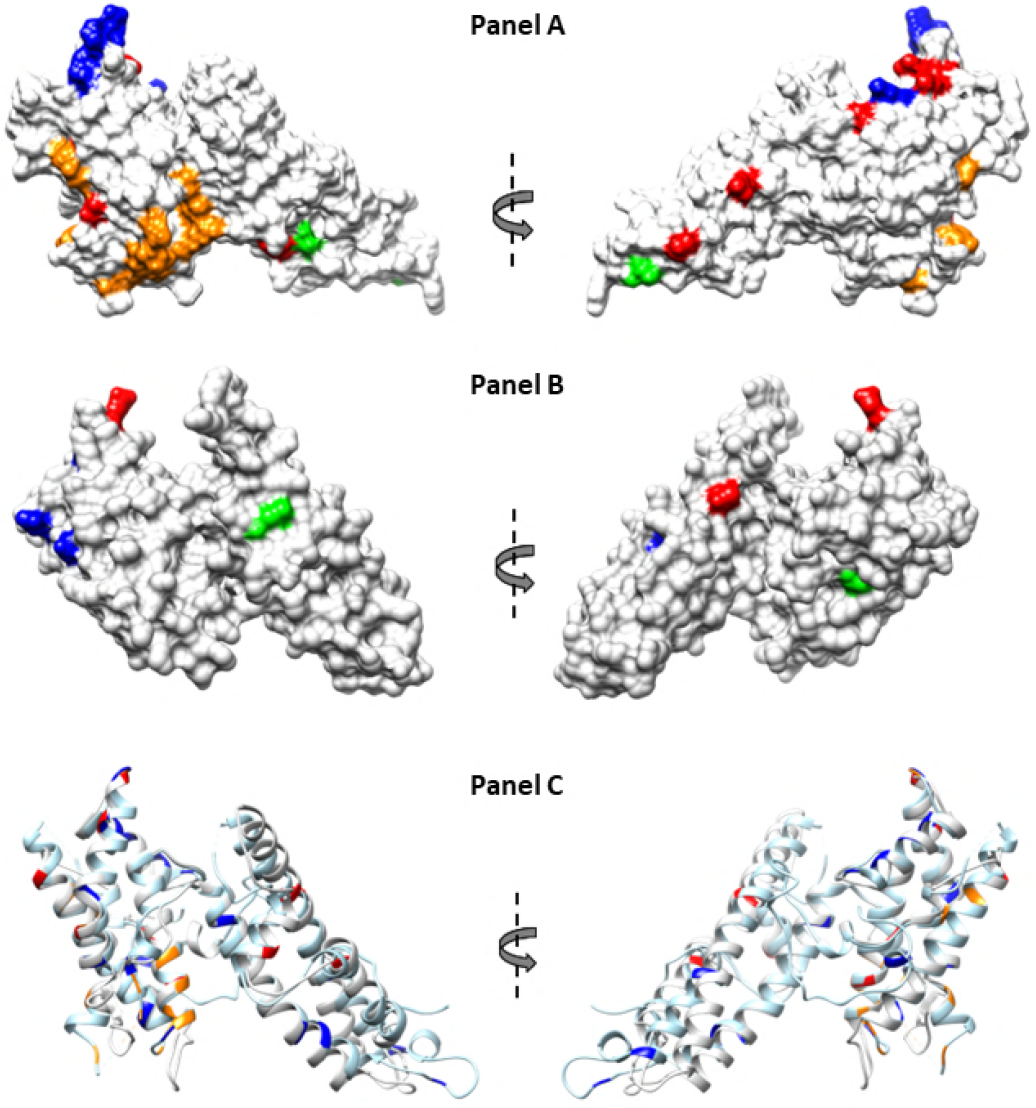
Mapping of SNPs on the structure of PvDBPII (Protein Data Bank code 4NUU) monomer and PvEBPII. SNPs present specifically to Cambodian and Madagascar isolates are highlighted in red and blue respectively, whereas SNPs present in the isolates from both the countries are highlighted in green. Putative binding residues on PvDBPII are highlighted in orange. Molecular surface diagram of PvDBPII is shown. SNPs and putative binding residues of PvDBPII are highlighted (Panel A). Predicted structure of PvEBPII is shown as molecular surface diagram and SNPs are highlighted (Panel B). Helical ribbon representation of PvDBPII (in light blue) and PvEBPII (in light grey) are superimposed. SNPs and putative binding residues are highlighted (Panel C).

A total of 169 *P. vivax* isolates with both SNP and CNV were available for analysis. Among them, 76 (45%) had multiple copies *PvDBP*. Gene amplification was observed in 9/18 alleles (50%) in Cambodia and in 8/9 alleles (89%) in Madagascar isolates (Fig 4, Panel A). No specific *PvDBP* allele was found to have in proportion more isolates than expected with *PvDBP* gene amplification, indicating that *PvDBP* gene amplification can occur across multiple alleles.

**Fig 4.**
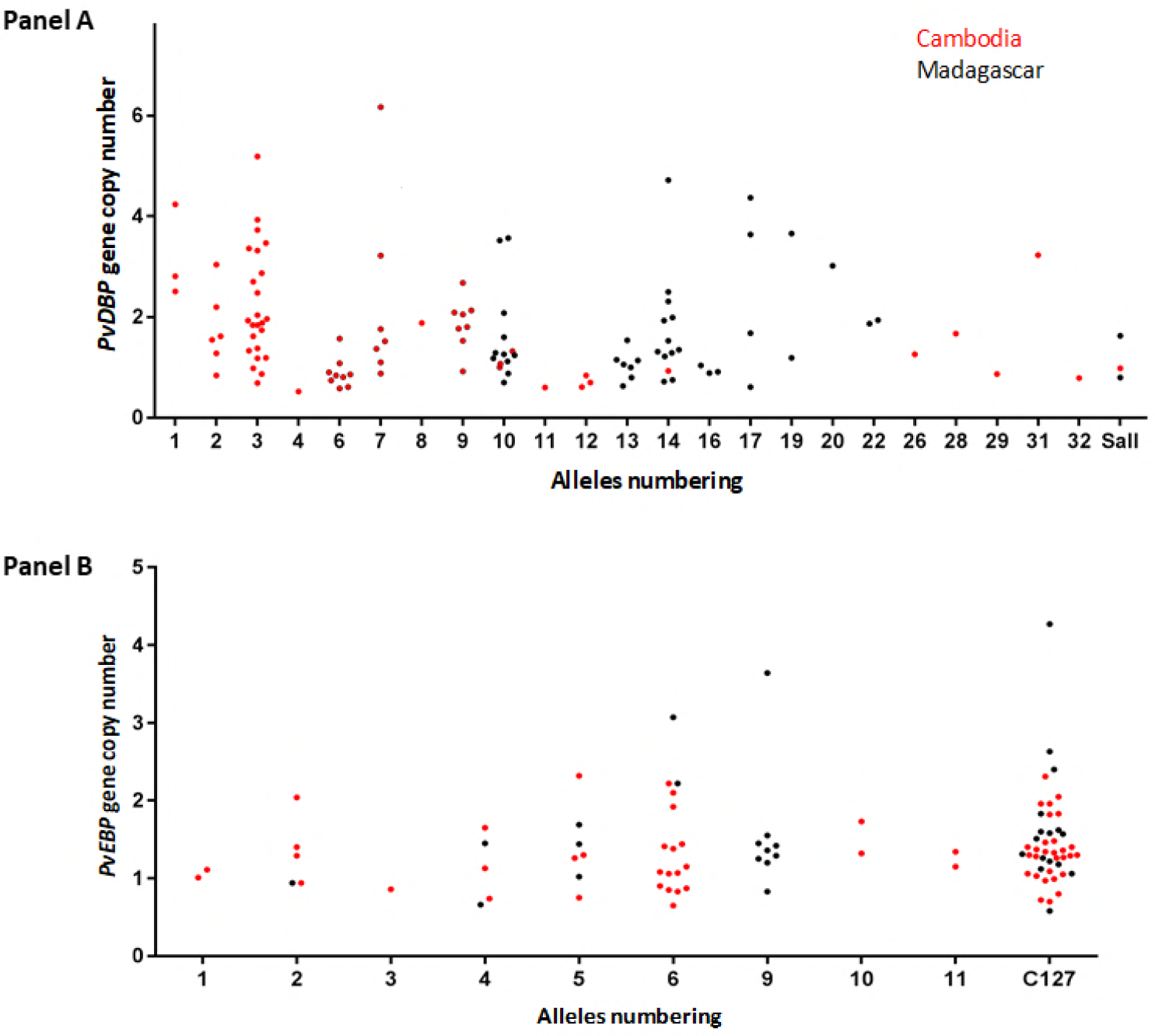
Distribution of *PvDBP* gene copy number of 33 PvDBP alleles in isolates from Cambodia and Madagascar (Panel A). Distribution of PvEBP gene copy number of 12 PvEBP alleles in isolates from Cambodia and Madagascar (Panel B).

### *PvEBP* gene copy number variation (CNV)

*PvEBP* CNV was determined in 184 Cambodian and 66 Malagasy samples (Table 1). *PvEBP* CNV ranged from 1 to 2 copies in Cambodian isolates and 1 to 5 copies in Malagasy samples. The proportion of multiple copies *PvEBP* isolates was significantly more frequent in Madagascar compared to Cambodia: 56% (37/66)*vs.* 19% (35/184) (P<10^−6^, Fisher’s exact test). The median copy number of *PvEBP* gene was also significantly higher in isolates from Madagascar compared to Cambodia: 1.6 (IQR: 1.3-2.1) *vs*. 1.3 (IQR : 0.95-1.4) (P<10^−4^, Mann Whitney test) (Fig 1, Panel B).

In Cambodia, the proportion of isolates with multiple copies of *PvEBP* or the median copy number of *PvEBP* gene was higher in high transmission areas located in eastern provinces (Ratanakiri, Mondulkiri and Kratie): 24% (31/127) vs. 7% (4/57) (P = 0.004, Fisher’s exact test), respectively and 1.3 (IQR: 1.0 - 1.5) *vs.* 1.1 (IQR: 0.85 - 1.3) (P = 0.003, Mann Whitney test). *PvEBP* CNV (proportion of isolates carrying *PvEBP* amplification or median copy of *PvEBP* gene) were similar between paired isolates collected at time of chloroquine treatment (D0) and on day of recurrence (Dx): D0, 60% (6/10), median *PvEBP* copy number= 1.6 (IQR: 1.4-1.7) vs. Dx, 50% (2/4), median *PvEBP* copy number=1.7 (IQR: 1.0-2.0) (P=1.0, Fisher’s exact test and P = 0.83, Mann Whitney test, respectively).

In Malagasy samples, no significant differences in the proportion of isolates with *PvEBP* amplification or the median copy of *PvEBP* gene were observed between homozygous (T/T) and heterozygous (T/C) Duffy-positives individuals: T/T, 54% (12/22), median *PvEBP* copy number=1.5 (IQR: 1.2-2.2) vs. T/C, 41% (12/29), median *PvEBP* copy number=1.4 (IQR: 1.2-1.7) (P=0.4, Fisher’s exact test and P=0.3, Mann Whitney test, respectively).

### *PvEBP* gene amplification occurs across multiple limited PvEBP alleles

*PvEBP* sequences obtained from 150 Cambodian and 69 Malagasy isolates (Table 1) were compared to the reference genome (C127) [20]. Eleven non-synonymous SNPs were observed leading to twelve different alleles. Most of them (59%) were the C127-like allele (Fig 5 and S2 Table).

**Fig 5.**
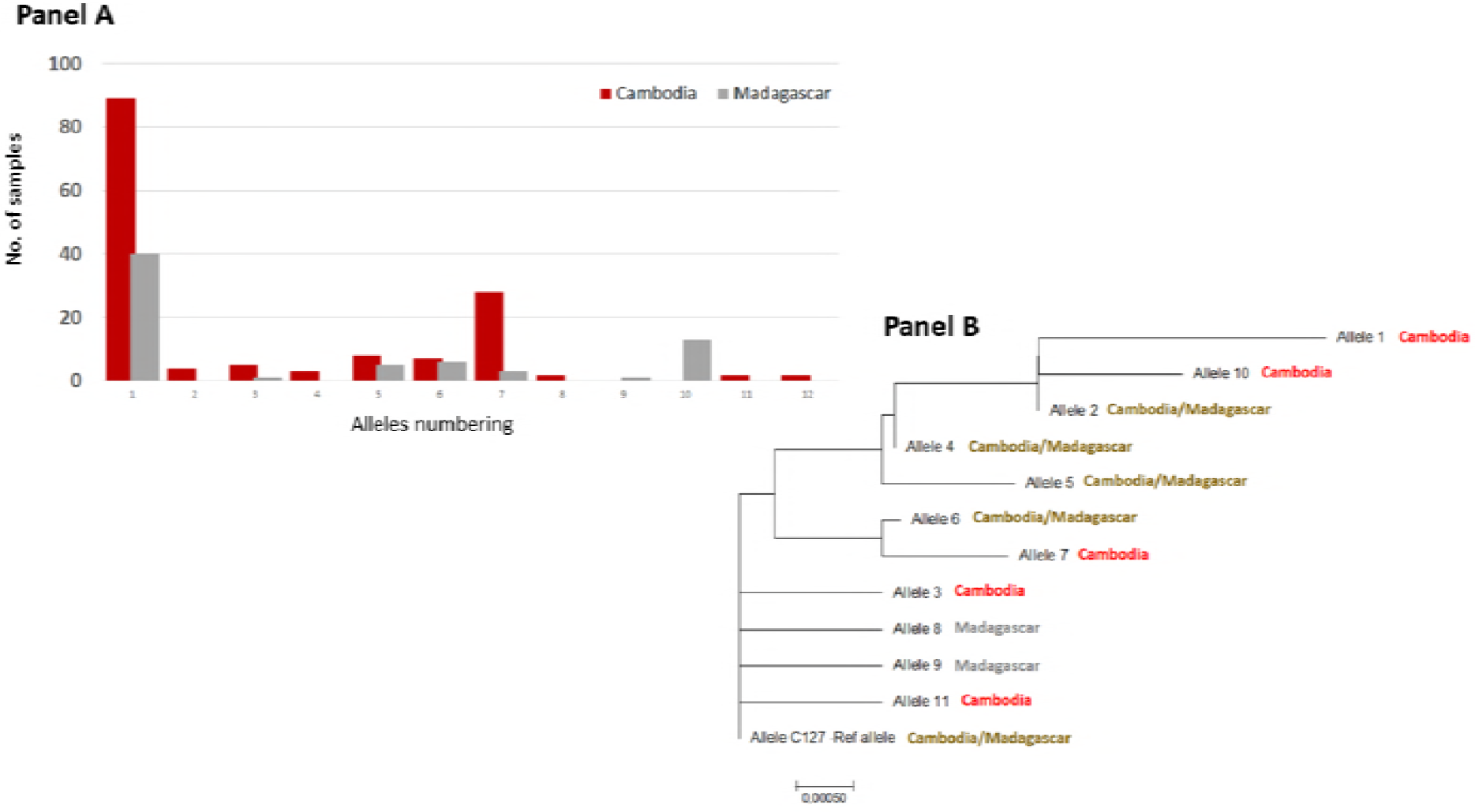
Distribution and number of *PvEBP* alleles among Malagasy and Cambodian samples (Panel A). Phylogenetic tree inferred using the Neighbor-Joining method (Panel B). Each allele was numbered 1-11 (see S2 Table). The evolutionary distances were computed using the Poisson correction method [23] and are in the units of the number of amino acid substitutions per site. The analysis involved 12 amino acid sequences. Evolutionary analyses were conducted in MEGA7 [24].

Four SNPs were common to both countries (N233K, D311N, E322K and I463V), three specific to Madagascar (N200K, C219S and N449K) and four to Cambodia (D268N, E306K, T421I, S423W). More common alleles were observed between the two countries for *PvEBP* compared to *PvDBP*: half of PvEBP alleles were shared between both countries compared to 1/11 for PvDBP alleles (P=0.02). Given that PvEBPII shares homology with DBL domains, the structure of PvEBPII was modeled based on the structure of PvDBPII (Fig 3, Panel B). The ribbon diagrams for the structures of PvDBPII and PvEBPII were superimposed and found to overlap (Fig 3, Panel C). SNPs were mapped to the PvEBPII structure and were found to be distributed across the domain. Since the receptor-binding site of PvEBPII remains to be defined, we were not able to assess whether the binding site was conserved or variable and deduce if this protein ligand can be targeted by strain transcending inhibitory antibodies to block receptor binding and invasion by diverse strains.

A total of 143 *P. vivax* isolates with both SNP and CNV data for *PvEBP* were available for analysis. Among them, 28 (20%) were found to have *PvEBP* gene amplification. *PvEBP* gene amplification was observed in 6/10 alleles (60%) in Cambodia and in 5/7 alleles (71%) in Madagascar isolates (Fig 4, Panel B). No specific *PvEBP* allele was found to have in proportion more isolates than expected with *PvEBP* gene amplification, indicating that *PvEBP* gene amplification can occur across multiple alleles.

## Discussion

The work presented here was focused to explore the polymorphism and the copy number variation of two *P. vivax* protein ligands involved in invasion into reticulocytes [41]. Unsurprisingly, we observed that SNPs detected in the receptor-binding domain of PvDBPII were similar to those observed previously. The binding residues for DARC in PvDBPII were conserved and majority of the SNPs observed were distal to the binding site. In fact, many SNPs were found on the opposite face of PvDBPII compared to the binding site suggesting that this surface is under immune pressure during natural infection. Genome sequencing of different *P. vivax* isolates collected in various malaria endemic settings had previously revealed that the PvDBP gene is frequently duplicated [17, 18]. Here, we confirm again that amplification of *PvDBPII* is a common genomic event, which can frequently occur in isolates from areas where Duffy-negative phenotype is virtually absent such as Cambodia. To encompass the two types of duplication previously observed [17, 18], we designed a novel quantitative real-time PCR with a standard curve that enables precise quantification of the number of PvDBP and PvEBP genes up to 6 copies. Our data confirms the hypothesis that *PvDBP* amplification is unlikely to play a role in invasion of Duffy-negative reticulocytes or at least that human Duffy-negative populations do not specifically select for parasites with multiple copies of gene encoding *PvDBP*.

The critical role of PvDBP protein ligand in invasion pathway into human Duffy-positive reticulocytes is well established while alternative pathways through other parasite protein ligands involved in the invasion of Duffy-negative reticulocytes are unknown. To date, few biological investigations seem to support a possible role of the newly described PvEBP protein ligand in the invasion of Duffy-negative reticulocytes [19, 21, 41]. By exploring its genetic diversity, we discovered two interesting findings. First, a high proportion of isolates from Madagascar (56%) where Duffy negative and positive individuals coexist had multiple copies of *PvEBP* compared to Cambodia (19%) where Duffy-negative population is virtually absent. Second, there was evidence for an excess of nonsynonymous mutations and the total absence of synonymous mutations in PvEBP. Indeed, among the 219 *P. vivax* tested samples we observed only eleven non-synonymous point mutations in PvEBP sequences and twelve different alleles (S2 Table). SNPs found were distributed across PvEBPII but we were not able to define whether polymorphism affects the binding site, which remains to be identified. In any case, the absence of synonymous mutations clearly reveals that this gene is under strong diversifying selection, as has been shown previously for *P. falciparum EBA-175* and *PvDBP* [42]. The most likely agent driving the diversification of these antigens is the human acquired immune response. Indeed, we can speculate that novel alleles encoding parasite antigens occuring in the population would potentially be able to avoid the human immune response and as such give the parasite a survival advantage leading to the allele’s selection. However, it remains unclear whether the elevated frequency of C127 allele in the population sample (~62% and ~52% in Cambodia and Madagascar, respectively) indicates a relatively recent selective increase of a new variant whose sequence may reflect its ability to avoid immune detection.

Due to a similar type of selection on *P. falciparum* EBA-175 and *P. vivax* EBP, our data confirms the importance of the newly decribed protein ligand PvEBP in invasion of human reticulocytes and as a target of acquired immunity. Furthermore, the role of PvEBP as a protein ligand is supported by recent serological analysis^41,42^. These studies have shown that antibodies against the region II domain of the PvEBP were commonly detected in sera from individuals in malaria endemic settings such as in Cambodia or in Solomon Islands/Papua New Guinea. In Cambodia, humoral immune response to PvEBP was found to be higher and longer lasting compared to PvDBP, making this marker a better candidate to monitor *P. vivax* infections [43]. Moreover, Franca *et al*, in Papua New Guinea identified a significant association between reduced risk of clinical vivax malaria and levels of antibodies against PvEBP [44].

As for PvDBP, we also assessed the *PvEBP* gene copy number in our array of isolates and observed that contrary to PvDBP, isolates from Madagascar, where Duffy negative and positive individuals coexist, carry more frequently a PvEBP gene expansion. In particular, we detected in Malagasy isolates a bimodal distribution of parasites with multiple copies of PvEBP that includes a specific population of parasites with > 3 copies of PvEBP gene that was not found in Cambodian isolates. Unfortunately, we could not test for association between the *PvEBP* gene expansion and their capacity to invade Duffy-negative reticulocytes, as reported recently by Gunalan *et al.* for PvDBP [19]. Indeed, among the few *P. vivax* samples collected from Duffy-negative individuals we had, we were not able to generate reliable PCR signals, probably because of the very small amount of DNA reflecting the usually low parasitemia found in these individuals.

One of the limitations in this work is that the majority of patients likely carry multiple clones of *P. vivax* [45, 46]. By using a qPCR approach as we did, we are aware that the PvDBP and PvEBP estimated gene copy number reflects the major clone contained in each isolate Next generation sequencing such as single cell DNA sequencing or even droplet PCR approach should be able to overcome this issue and assess CNV for each clone. Similarly, the Sanger sequencing technology used in this work to determine the sequences of PvDBP and PvEBP does allow sequencing only the major clone, preventing us to determine whether each strain in isolates with multiple copies of PvDBP and PvEBP had similar or different alleles.

In summary, we provide here data regarding the genetic diversity of the *PvEBP* gene, a recently described and potential protein ligand involved in invasion into human reticulocytes. Evidence for positive diversifying selection on the region II domain of the PvEBP was observed, which is similar to the evidence for diversifying selection on PvDBP region II. These observations clearly confirm the importance of PvEBP in the invasion process of the human reticulocyes and/or as a target of acquired immunity. In addition, the high proportion of *P. vivax* isolates with multiple copies of PvEBP gene that we found in Madagascar is intriguing and needs additional in-depth investigations. So far, the association between invasion in Duffy-negative individuals and *P. vivax* specific genomic traits (SNP or CNV) mainly relies on epidemiological associations. Direct evidence through reticulocyte invasion assays *in vitro* or in humanized mouse models with *P. vivax* isolates with diverse alleles and copy numbers of PvDBP and PvEBP and human reticulocytes with different Duffy phenotypes will be of great value in deciphering the molecular basis of Duffy-independent invasion pathway(s) used by *P. vivax*.

## Materials and Methods

### Study sites and samples collection

One hundred twenty nine Malagasy *P. vivax* samples were collected in 2015-2017 through cross-sectional surveys in the district of Maevatanana, from asymptomatic individuals during active case detection and from symptomatic patients seeking antimalarial treatment in health centers located in three communes (Andriba, Antanimbary and Maevatanana). *P. vivax* infections were detected using a malaria rapid diagnostic test (CareStart^TM^ Malaria Pf/pan RDTs, Accesbio) and, capillary blood samples were spotted into filter papers for each positive case.

In Cambodia, 453 *P. vivax* isolates were collected in 2003-2017 from symptomatic patients, seeking antimalarial treatment in health centers located around the country. Malaria diagnosis was also performed by RDT (CareStart^TM^ Malaria Pf/pan RDTs, Accesbio) and microscopy. Venous blood samples were collected from confirmed vivax malaria cases into 5 ml EDTA tubes. We also included in our analysis, 16 *P. vivax* isolates collected from recurrences occurring in the 2-months follow up after a standard 3-day course of chloroquine (30 mg/kg) in patients relocated in a non-transmission area (Popovici *et al.,* submitted manuscript).

The study protocols were reviewed and approved by the Cambodian National Ethics Committee on Health Research (IRB 038NECHR) or the National Ethics Committee in Madagascar (Ministry of Health, 141/MSANP/CE). All individuals or their parents/guardians provided informed written consent before sample collection.

### DNA extraction and PCR confirmation of vivax malaria

DNA was extracted from blood spots with Instagene Matrix reagent (BioRad, Marnes-la-Coquette, France) or from whole blood samples using the QIAamp DNA Blood Mini Kit (Qiagen, Courtaboeuf, France), according to the manufacturer’s instructions. Molecular detection and identification of *Plasmodium* parasites were performed by using real-time PCR targeting the *cytochrome b* gene as described previously [47, 48].

### SNPs analysis in *PvDBP* and *PvEBP* genes

*PvDBP* and *PvEBP* sequences were determined by nested PCR targeting the *PvDBP* region II and Sanger sequencing (Macrogen, Seoul, South Korea) using the following conditions. A first round PCR was conducted in 25 μL reactions using DNA, 0.2 μM of primers, 250 μM each dNTP, 2 mM MgCl2, and 1.25 units Taq Solis DNA Polymerase (Solis BioDyne, Tartu, Estonia) under the following conditions: 94 °C for 15 min, followed by 40 cycles of 94 °C for 20 s, 56 °C for 40 s, 72 °C for 90 s, and a final extension at 72 °C for 10 min. The nested PCR was carried out in 55 μL reactions using 2 μL of the primary PCR products diluted at 1/10, 0.40 μM of primers, 250 μM each dNTP, 2.5 mM MgCl_2_, and 2.5 units Taq Solis DNA Polymerase (Solis BioDyne, Tartu, Estonia) under the following conditions: 94°C for 15 min, followed by 40 cycles of 94 °C for 20 s, 60 °C for 20 s, 72 °C for 60 s, and a final extension at 72 °C for 10 min (S3 Table). Nucleotides and corresponding amino acids were analyzed using the CEQ 2000 software (Beckman). The sequences generated were compared to PVX_110810 (Pv_Sal1_chr06:976,329-980,090 (+)) for PvDBP and to *P. vivax* isolate C127 nEBP gene (KC987954.1) for PvEBP.

### CNVs analysis of PvDBP and PvEBP genes

*PvDBP* and *PvEBP* genes copy numbers were measured relatively to the single copy β-tubulin gene (housekeeping gene) using a CFX96 real-time PCR thermocycler (Biorad, Singapore). PCR were performed in 20 μL volumes in a 96-well plate containing 1X HOT FIREPol EvaGreen qPCR Mix Plus (Solis BioDyne, Tartu, Estonia), 0.5 μM of each forward and reverse primer and 2 μL of template DNA. Amplifications were performed under the following conditions: 95°C for 15 min, followed by 45 cycles of 95°C for 15 s, 60°C for 20 s, and 72°C for 20 s. *PvDBP* and *PvEBP* genes copy numbers were estimated in triplicate relative to a standard curves by using synthetic genes cloned in pEX-A2 vector (Eurofins Genomics, Greece) (*β-tubulin, PvDBP, PvEBP*) mixed at different ratio from 1:1 up to 1:11 (1 copy of (β-tubulin and 1 to 6 copies of *PvDBP* or *PvEBP*) (S3 Table). The ΔCT method (where CT is the cycle threshold) was used to determine the number of copies of each sample. In addition, an isolate with one copy of each gene was used as control. All isolates with copy number estimates of less than 0.5 were discarded.

### Duffy genotyping

Duffy genotypes were determined by nested PCR and Sanger sequencing (Macrogen, Seoul, South Korea) as previously described [49]. The inner PCR was conducted in 25 μL reactions using 3 μL of template DNA, 0.4 μM of primers, 250 μM each dNTP, 2 mM MgCl_2_, and 1.25 units Taq Solis DNA Polymerase under the following conditions: 94 °C for 15 min, followed by 40 cycles of 94 °C for 30 s, 58 °C for 30 s, 72 °C for 90 s, and a final extension at 72 °C for 10 min. Outer PCRs detecting mutations in the GATA box (T or C) and in the coding sequence were carried out in 55 μL reactions using 2 μL of the primary PCR products diluted at 1/10, 0.36 μM of each primer), 250 μM each dNTP, 2.5 mM MgCl_2_, and 1.25 units Taq Solis DNA Polymerase under the following conditions: 94 °C for 15 min, followed by 40 cycles of 94 °C for 20 s, 58 °C for 20 s, 72 °C for 60 s, and a final extension at 72 °C for 10 min (S3 Table).

### Mapping SNPs on structural models of PvDBPII & PvEBPII

Coordinates of single chain of PvDBP (PDB ID 4NUU) were obtained from Protein databank [50, 51] and used for mapping SNPs. SNPs were mapped on the PvDBPII structure using Chimera software [52]. In addition, putative binding residues of PvDBPII predicted previously [38–40] were also mapped on the PvDBPII structure.

The selected primary PvEBPII sequence (allele C127) [20] (Figure S1) was used for 3D modeling prediction (S1 Figure). 3D Model of the PvEBPII domain was determined by homology based structure prediction online tool using Phyre2 under default mode [53]. 3D structure with maximum score was selected and highly flexible N- and C-terminal ends were truncated from the structure. Refinement of predicted 3D model and minimization of local structural distortions was performed using ModRefiner [54]. The overall quality of the predicted 3D model was evaluated with QMEANDisCo (Qualitative Model Energy ANalysis- Distance Constraint) score. QMEANDisCo is a tool for assessing the agreement of pairwise residue-residue distances with ensembles of distance constraints extracted from structures homologous to the assessed model [55]. SNPs were mapped on the PvEBPII model using Chimera software [52].

### Statistical analysis

Data were analyzed with Microsoft Excel and MedCalc version 12 (Mariakerke, Belgium). Quantitative and qualitative data were expressed as median (IQR) or proportion (%), respectively. The Mann-Whitney *U* test was used for non-parametric comparisons. For categorical variables, proportions were examined by Chi-squared or by Fisher’s exact tests. Two-sided p-values of <0.05 were considered statistically significant.

MUSCLE multiple alignment and evolutionary analyses were conducted in MEGA [56] with 1000 bootstrap replicates. All positions containing gaps and missing data were eliminated. Phylogenetic reconstructions were performed using neighbor joining (NJ).

## Acknowledgements

We acknowledge all patients from Cambodia and Madagascar participating to the study along with health care staff located in health centers.

## Author contributions statement

C.R., J.P., S.R., M.G.B., C.H., D.S., C.C., I.V.W., D.M. conceived the experiments, S.K., S.B., V.R. and E.R., R.L.R., T.R., Z.A. conducted the sample collection in Cambodia and Madagascar, respectively. Molecular analysis were performed by C.R., S.B., V.R. at Institut Pasteur in Cambodia and by E.R., R.L.R., T.R., Z.A. at Institut Pasteur in Madagascar. A.K. and C.C. did the mapping SNPs on structural models of PvDBPII & PvEBPII. C.R., J.P., S.R., I.V.W., and D.M. conducted the statistical analysis. C.R., J.P., C.C., I.V.W. and D.M. analysed the results. C.R., J.P., C.C., I.V.W. and D.M.wrote the draft manuscript. All authors reviewed the manuscript.

## Supporting information

**S1 Table. List of SNP and alleles observed in *P. vivax* Duffy binding sequences in isolates collected from Cambodia (N=153) and Madagascar (N=92).** PVX_110810, Pv_Sal1_chr06:976,329-980,090 (+) was used as reference.

**S2 Table. List of SNP and alleles observed in *Plasmodium vivax* erythrocyte binding sequences in isolates collected from Cambodia (N=150) and Madagascar (N=69).** *P. vivax* isolate C127 nEBP gene, KC987954.1 was used as reference.

**S3 Table. List of the primers and synthetic genes along with their nucleotide sequences used to assess the genetic diversity (SNP and CNV) of PvDBP and PvEBP and for Duffy gentotyping.**

**S1 Figure. *PvEBP* reference sequence (allele C127) used to predict the 3D stucture (middle portion presente in red font was obtained by PCR/sequencing and the sequences sequences located at the ends was deduced from PVX_110810, Pv_Sal1_chr06:976,329-980,090).**

